# Newton’s Cradle: Cell Cycle Regulation by Two Mutually Inhibitory Oscillators

**DOI:** 10.1101/2024.05.18.594803

**Authors:** Calin-Mihai Dragoi, John J. Tyson, Béla Novák

**Affiliations:** Department of Biochemistry, University of Oxford, South Parks Road, Oxford OX1 3QU, UK; Department of Biological Sciences, Virginia Tech, Blacksburg, VA 24061, USA

**Keywords:** cell cycle, bistability, oscillations, checkpoints, endocycles

## Abstract

The cell division cycle is a fundamental physiological process displaying a great degree of plasticity during the course of multicellular development. This plasticity is evident in the transition from rapid and stringently-timed divisions of the early embryo to subsequent size-controlled mitotic cycles. Later in development, cells may pause and restart cell proliferation in response to myriads of internal or external signals, or permanently exit the cell cycle following terminal differentiation or senescence. Beyond this, cells can undergo modified cell division variants, such as endoreplication, which increases their ploidy, or meiosis, which reduces their ploidy. This wealth of behaviours has led to numerous conceptual analogies intended as frameworks for understanding the proliferative program. Here, we aim to unify these mechanisms under one dynamical paradigm. To this end, we take a control theoretical approach to frame the cell cycle as a pair of arrestable and mutually-inhibiting, doubly amplified, negative feedback oscillators controlling chromosome replication and segregation events, respectively. Under appropriate conditions, this framework can reproduce fixed-period oscillations, checkpoint arrests of variable duration, and endocycles. Subsequently, we use phase plane and bifurcation analysis to explain the dynamical basis of these properties. Then, using a physiologically realistic, biochemical model, we show that the very same regulatory structure underpins the diverse functions of the cell cycle control network. We conclude that Newton’s cradle may be a suitable mechanical analogy of how the cell cycle is regulated.

**Declaration of interest:** The authors declare no competing or financial interests.

## Introduction

The cell cycle is a recurrent sequence of events whereby a eukaryotic cell accurately replicates all its chromosomes during S phase (DNA synthesis) and then partitions the replicated chromosomes (‘sister chromatids’) evenly to two daughter cells during M phase (mitosis and cell division), so that each daughter cell receives a full and faithful copy of the organism’s genome (Fig. 1). For this process to maintain genetic continuity from one generation of cells to the next, it is essential that the processes of DNA replication and mitosis alternate, and that any damage to the DNA sequences or to chromosome integrity be repaired before the DNA is replicated or the sister chromatids are pulled apart on the mitotic spindle [1].

**Figure 1.**
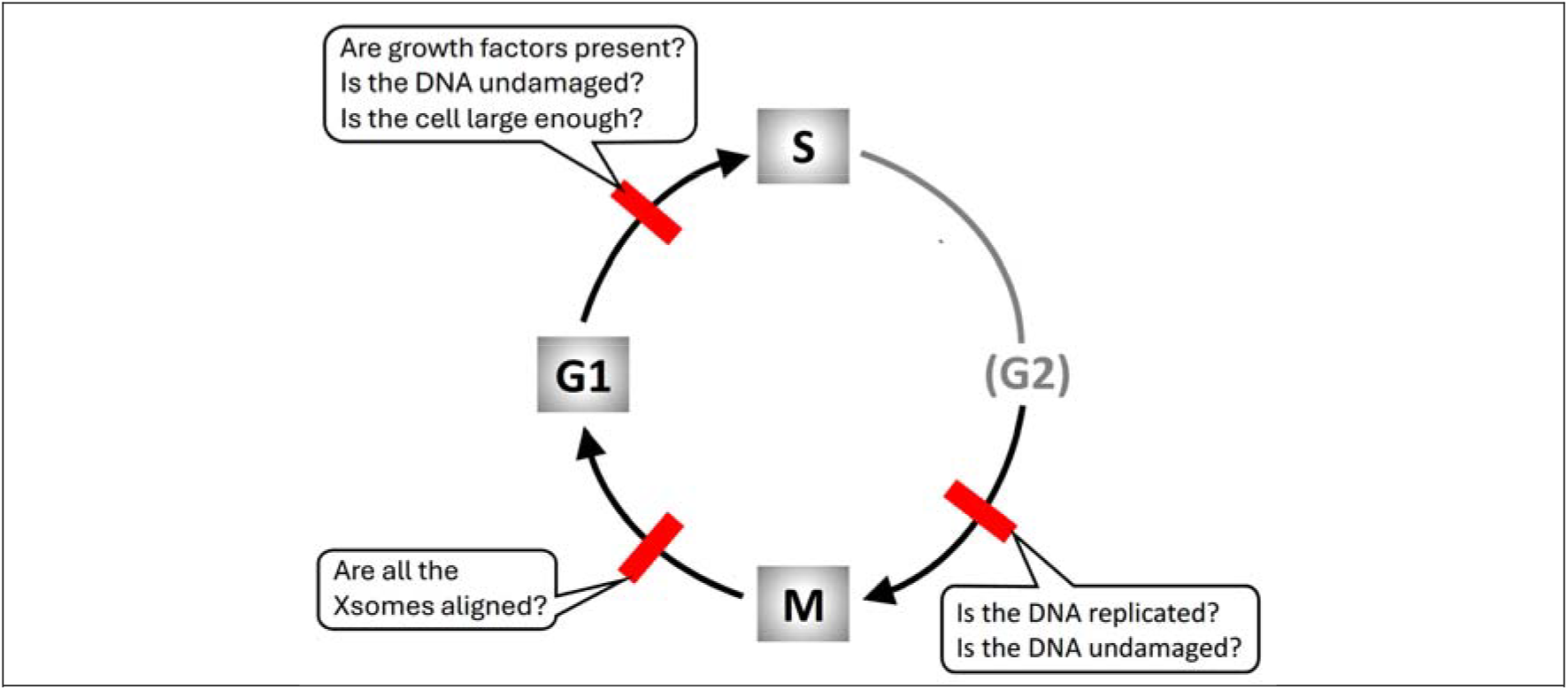
The eukaryotic cell cycle. Cell reproduction consists of a phase of DNA synthesis (S phase), when every chromosome is replicated, followed in time by the process of mitosis (M phase), when the replicated chromosomes are segregated to opposite poles of the mitotic spindle and the mother cell divides into two daughter cells, each containing a complete set of (unreplicated) chromosomes (i.e., G1 cells). Successful cell proliferation requires strict alternation of S phase and M phase, which is assured by three ‘checkpoints’ (red bars) that block subsequent steps of the cycle until certain conditions are fulfilled. G2 = phase after completion of DNA replication and before entry into mitosis.

The integrity of genomic transmission during the eukaryotic cell cycle is guarded at three major ‘checkpoints.’ At the G1/S transition, the cell checks that it is exposed to the proper growth factors, that its DNA is undamaged, and that it is large enough to warrant a new round of DNA replication and division. At entry into mitosis, the cell checks that its DNA is fully replicated and undamaged. And before the cell attempts to exit from mitosis, it checks that all its sister chromatid pairs are properly aligned on the mitotic spindle.

### Metaphors for cell cycle regulation

In light of these basic facts about eukaryotic cell division, many different metaphors have been proposed to understand how the events of the cell cycle are controlled.

#### Dominoes

Early researchers viewed progression through the cell division cycle as a dependent sequence of events, like a line of falling dominoes [2]. From the simple sequence G1→S→G2→M→G1→…, there evolved more complex, mechanistic models based on hypothetical mitotic activators and inhibitors [3; 4]. These views were reinforced by genetic studies in yeasts [5; 6], which culminated in a Boolean model of the yeast cell-cycle control system [7] with one gene product flipping the next in a robust sequence of dependent molecular events.

#### Clock

Other early researchers, focussing on the periodicity of the cell division cycle, envisioned the control system as a clock, ticking away and sending out commands for DNA replication, mitosis and cell division [8; 9]. This view was reinforced by biochemical studies of the autonomous, periodic activation of Mitosis Promoting Factor (MPF) in frog egg embryos [10], culminating in limit-cycle models of MPF synthesis, activation and degradation [11; 12; 13].

#### Clock Shop

More recently, researchers have recognized that, when certain genes of the cell-cycle control system are mutated, then subsets of cell cycle events can repeat periodically; for example, rounds of DNA replication without mitosis [14], periodic budding of yeast cells without division [5; 15], periodic fluctuations of metabolism [16; 17], transcription factors [18], and other proteins [19; 20] without DNA synthesis or mitosis. These observations led to the notion of the cell-cycle control system as a collection of oscillators whose interactions govern both normal cell cycle progression and the periodicities observed when the G1→S→G2→M sequence is short-circuited or arrested. The clock-shop hypothesis is often modelled by Kuramoto’s equations [21] for a population of phase-coupled oscillators, *dθ*_*i*_/*dt* = *ω*_*i*_ + ∑_*j≠i*_ *k*_*ij*_ sin (*θ*_*j*_ - *θ*_*i*_).

#### Copy Machine

First and foremost, the mitotic cell cycle is a ‘copy machine’ that makes a copy of each chromosome and then collates a complete set of chromosomes to each of two daughter cells. Like the office copy machine, the cell cycle can spit out daughter cells with clock-like regularity if demand for its services is high, or only irregularly if demand is low. Like the cell cycle, the office copy machine is guarded by ‘checkpoints’: out-of-paper, out-of-toner, out-of-staples, and the dreaded ‘paper jam.’ Like the service technician who needs to know the internal mechanisms of the copy machine, the cell biologist needs to know the molecular mechanisms of DNA replication and mitotic spindle function and the molecular controls that govern these processes and their checkpoints. Models of these mechanisms are usually formulated as sets of nonlinear differential equations [22; 23].

#### Toggle Switches

It is commonplace now to think of checkpoint controls as toggle switches that can be flipped between stable OFF and ON states [24; 25]. Before the transition, the switch is OFF, because inhibition signals outweigh activation signals. When activation prevails, the switch flips ON, the transition proceeds, and the switch stays in the ON position until later events of the cell cycle restore the switch to OFF. The toggle switch metaphor accounts nicely for the ‘irreversibility’ of cell cycle progression and the roles of checkpoints guarding genome integrity. The latest incarnation of this view is the ‘latching gates’ hypothesis of Novak & Tyson [26; 27], which provides a unified description of both normal cell cycle progression and of ‘endocycles’ (G1→S→G1→S and M→G1→M→G1).

In this paper we would like to elaborate on the latching-gates model to reconcile the clock-shop and toggle-switch views of cell cycle control. Along the way, we introduce a new metaphor—Newton’s Cradle—that may provide a useful way to think about the cell cycle control system.

### Mechanisms of cell cycle regulation

The eukaryotic cell division cycle is controlled by a complex network of interacting genes and proteins, focussed on regulating the activities of a family of protein kinases called cyclin-dependent kinases (CDKs). During S-(G2)-M phases of the cell cycle, CDK activities are high, because CDKs are crucial drivers of DNA synthesis and early mitotic events. As a cell exits mitosis and divides, all CDK activities are extinguished, and, consequently, the daughter cells are born in G1 phase with low CDK activity. At the G1/S transition, CDK activity is abruptly regained [1].

Figure 2 summarizes the molecular interactions governing CDK activation and inactivation during the cell cycle [27]. The diagram, though incomplete to be sure, is sufficient for our purposes. The most prominent components of the control network are the S-CDK (Cdk2:CycA) and the M-CDK (Cdk1:CycB) that drive DNA synthesis and mitosis, respectively. Each CDK is a heterodimer of a cyclin subunit (Cyclin A or Cyclin B) and a kinase subunit (Cdk2 or Cdk1). These CDKs have a mutual antagonistic relationship with the cyclin-degradation machinery, called the Cyclosome (also known as the Anaphase Promoting Complex; hence, APC/C). The antagonistic relationship between CDKs and the APC/C lies at the heart of cell cycle control [28] by creating a bistable switch with two qualitatively different states [27]. In G1 phase, APC/C:Cdh1 activity is high and both S-CDK and M-CDK activities are low. If internal and external conditions favour proliferation, the E2F transcription factor is activated [29], inducing synthesis of cyclins E and A. Rising activity of Cdk2:CycE phosphorylates Cdh1, causing it to dissociate from APC/C. Now Cdk2:CycA (S-CDK activity) can rise and initiate DNA replication [30]. In addition, Cdk2:CycA induces the synthesis of Cyclin B and rising activity of M-CDK. Both S- and M-CDKs keep Cdh1 phosphorylated and inactive.

**Figure 2.**
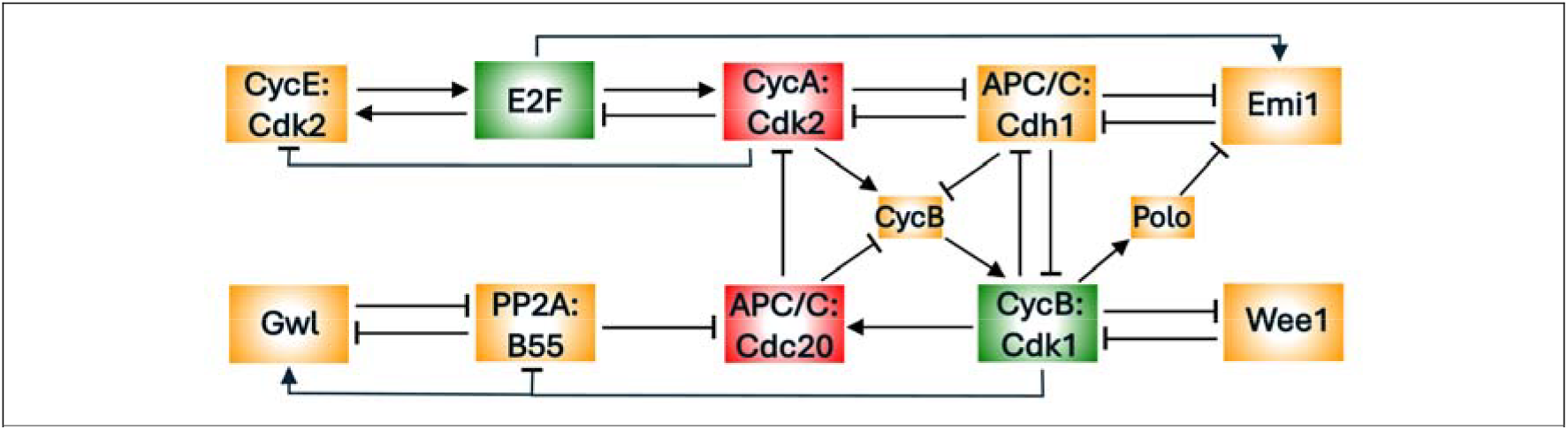
A simplified diagram of the cell-cycle regulatory network (from Dragoi et al., 2024). Cdk2:CycA and Cdk1:CycB are the cyclin-dependent kinases that initiate DNA synthesis and mitosis, respectively. They are both involved in double-negative feedback loops with the ubiquitin ligase, APC/C:Cdh1, creating two bistable switches that control the transitions from G1 to S and from M back to G1. The network has many other positive feedback loops (+ + or − −) and negative feedback loops (+ −, + + −, etc.) that are conducive to bistable switches and periodic oscillations.

Positive (and double-negative) feedback motifs within the network give rise to a hierarchy of bistable toggle switches that organise the proliferative program into clearly defined phases, during which specific physiological processes are carried out [31; 32]. The negative feedback loops within the network enable the switches to reset as the cell proceeds from one phase to the next; and they also set up the possibility for autonomous oscillations in subsets of cell cycle events. This summary of the cell-cycle control system suggests that we might envision it as two modules: an S-module (the upper row of Fig. 2) and an M-module (the lower row of Fig. 2) that have the capacity to oscillate (in isolation) but interact by mutual inhibition so as to flip back and forth between DNA synthesis and mitosis.

This view of the control system gave rise to our latching-gate description of cell cycle progression [26; 27]. In this scenario, normal mitotic cycles can be viewed as a strict alternation of G1 phase (APC/C:Cdh1 activity high, CDK activity low) and S-(G2)-M phase (APC/C:Cdh1 activity low, CDK activity high). The bistable switch is flipped from G1 into S by the transient activation of Cdk2:CycE, and from M back to G1 by the transient activation of APC/C:Cdc20. The model predicts that perturbations that render the bistable switch reversible may allow transitions back-and-forth between G1 and S or between M and G1.

Such oscillations, which span only a subset of the canonical phases, are called ‘endocycles.’ For instance, inhibition of Cdk1:CycB could prevent mitotic events and render the G1/S transition reversible (i.e., unlatched) with respect to Cdk2:CycE. This could cause periodic activation of Cdk2:CycA, driving discrete S phases in the absence of mitosis. This not-uncommon phenomenon is known as endoreplication [33]. In contrast, raising the basal activity of Cdk1:CycB could cause constitutive phosphorylation and inhibition of APC/C:Cdh1, thereby preventing the cell from returning to G1. Nonetheless, the negative feedback loop in the M-module may render the M/G1 transition reversible with respect to APC/C:Cdc20, causing small amplitude oscillations in CycB concentration [27; 34].

By probing the state of the APC/C:Cdh1 – Cdk1:CycB switch, the latching-gate view provides a top-down perspective on the dynamical processes that distinguish mitotic cycles from endocycles. Nonetheless, the molecular biology summarized in Fig. 2 is so complex that it obscures the minimal regulatory properties necessary for maintaining robust, irreversible mitotic cycles under most circumstances while also permitting multiple oscillatory modes (*endocycles*) in response to specific perturbations. Our aim here is to identify the necessary and sufficient regulatory structures in Fig. 2 that underlie both normal mammalian mitotic cycles and out-of-the-ordinary endocycles. At the core of our model are two oscillators (representing the S- and M-modules) coupled by mutual inhibition.

First, we propose a ‘toy’ model of the double-oscillator hypothesis in terms of two symmetric nonlinear second-order oscillators coupled by mutual inhibition. We show that the emerging ‘activity handover’ between the two oscillators is sufficient to explain the alternation of S- and M-phase events observed during mitotic cycles. Subsequently, we show that permanent arrest of one oscillator may leave the other one intact, giving rise to endocycles. Lastly, we show that the double-oscillator hypothesis is perfectly consistent with the latching-gate view.

After analysing the toy model, we return to the full model (Fig. 2) and show, using pseudo-phase plane analysis and bifurcation diagrams, that the CDK—APC/C mechanism known to operate in mammalian cells displays all the same qualitative features as the minimal model of four ODEs.

## Results and Discussion

### Two nonlinear oscillators drive replicative and mitotic events, respectively

Our depiction of the eukaryotic cell cycle control system in Fig. 2 identifies two analogous subsystems, an S-phase control module and an M-phase control module, that regulate the activities of Cdk2:CycA and Cdk1:CycB, respectively. At the core of each module is a negative feedback loop (NFL): in the S-phase module, Cdk2:CycA inactivates its transcription factor, E2F [35], while in the M-phase module, Cdk1:CycB activates APC/C:Cdc20, which targets cyclin B for degradation [36]. An NFL, which is a necessary attribute of all biochemical oscillators [37], is composed of (at least) two components, which can be can be called the Activator (e.g., E2F) and the Inhibitor (e.g., Cdk2:CycA). Another common feature of the two modules is that, in each case, both the activator and the inhibitor are amplified by positive or double-negative feedback loops (PFL or DNFL). In the S-phase module, E2F (the activator) induces the synthesis of CycE, while CycE activates E2F by inactivating its repressor Rb (not shown). Similarly, Cdk2:CycA (the inhibitor) hyper-phosphorylates and inactivates its antagonist, APC/C:Cdh1. In the M-phase module, the activator Cdk1:CycB inhibits its inhibitory kinase, Wee1. On the other side, the inhibitor APC/C:Cdc20 is amplified by the DNFL between PP2A:B55 and Gwl [38], which can flip the inhibitory role of APC/C:Cdc20 between ON and OFF states. According to Novak & Tyson [37], these network motifs, termed activator-inhibitor amplified negative feedback oscillators, give rise to robust and highly tunable oscillations.

### A double-oscillator motif illustrates the endo-oscillatory characteristics of the cell cycle

#### The basic oscillator Ω

To begin building-up the double oscillator motif, we first design a double-amplified negative feedback oscillator, Ω_1_ (Fig. 3A). Strictly speaking, the double amplification is not required for oscillations: a single positive feedback loop per oscillator can yield qualitatively similar results. Nonetheless, in order to highlight the similarity of Ω_1_ to the cell cycle control system (Fig. 2), we elect to add positive feedback to both the Activator and the Inhibitor. Hence, the dynamical system Ω_1_ consists of two nonlinear differential equations:

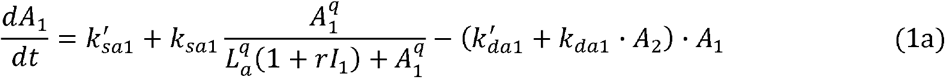

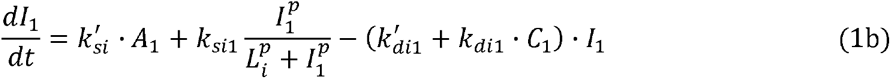

The synthesis of Activator_1_ (A_1_) is autocatalytic, according to a Hill function, and inhibited by Inhibitor_1_ (I_1_), which increases the effective binding constant, *L*_*a*_^*q*^ (1+*rI*_1_). The degradation of A_1_ is promoted by A_2_, the Activator of the other module (Ω_2_), which we take as a constant for now. The synthesis of Inhibitor_1_ (I_1_) is also autocatalytic, while its degradation is stimulated by a checkpoint component (C_1_).

**Figure 3.**
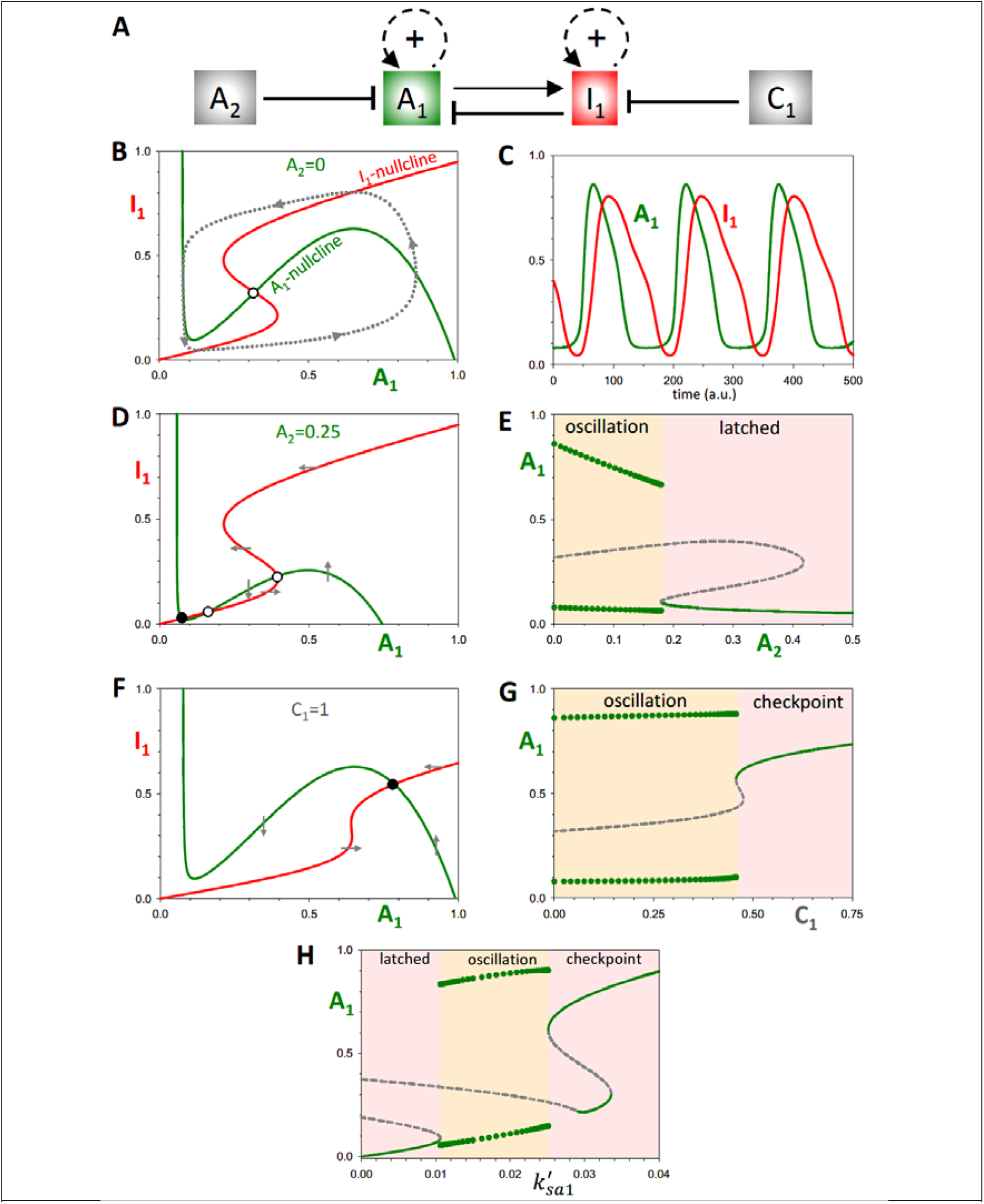
The dynamical properties of the single oscillator, Ω_1_. **(A)** Influence diagram of the single oscillator network. *A*_1_ and I_1_ are the activator and inhibitor of Ω_1_; *A*_2_ is the ‘latching’ parameter and *C*_1_ the ‘checkpoint’ parameter. **(B)** Phase plane of the Ω_1_ system in the basal oscillating state. Parameter values in the Material and Methods. The two nullclines intersect at an unstable steady state (o), surrounded by a limit cycle oscillation (dotted line). **(C)** Time course simulation of the limit cycle oscillation. **(D)** Phase plane for A_2_ = 0.25. The downward translation of the *A*_1_ nullcline results in three steady states: an unstable node (o), a saddle point (o), and a stable ‘latched’ state (●). The latched state arrests the oscillation at low activities of A_1_ and I_1_. **(E)** Bifurcation diagram of *A*_1_ with respect to the latching parameter, *A*_2_, showing the abrupt arrest of the oscillation at a SNIC bifurcation. **(F)** Phase plane for the checkpoint arrested system, when *C*_1_ = 1. The system arrests at a new stable steady state (●) with high *A*_1_ activity. **(G)** Bifurcation diagram of *A*_1_ with respect to the checkpoint parameter, *C*_1_, showing the abrupt arrest of the oscillation through a SNIC bifurcation. **(H)** Bifurcation diagram, *A*_1_ versus basal synthesis rate, *k’*_*sa1*_.

We start our analysis with an isolated system Ω_1_, when both A_2_ and C_1_ are set to zero. The phase plane of this system reveals a backwards N-shaped nullcline for A_1_ and an S-shaped nullcline for I_1_ (Fig. 3B). The two nullclines intersect at an unstable steady state, and the system exhibits limit cycle oscillations (Fig. 3B,C).

Next, we study the effect of increasing the value of A_2_. As shown in Fig. 3D, increasing *A*_2_ to 0.25 squashes down the A_1_-nullcline. This makes sense because both A_2_ and I_1_ inhibit A_1_, so, when A_2_ increases, a lower activity of I_1_ is needed to inactivate A_1_. In this case (*A*_2_ = 0.25), the nullclines intersect in three steady states, but only the leftmost steady state (marked with a black dot) is stable. We call this stable steady state the *latched state* and *A*_2_ the latching parameter. When the latched state exists, the dynamical system is attracted to it, suppressing oscillations and maintaining both A_1_ and I_1_ inactive.

To see how the system changes over a range of values of the latching parameter, we plot a bifurcation diagram (Fig. 3E), which shows the steady state value of A_1_ as a function of A_2_. The plot reveals that for sufficiently small values of *A*_2_, *A*_1_ oscillates with large amplitude. As *A*_2_ increases, oscillations arrest abruptly due to the appearance of the latched state at a SNIC (*saddle node on invariant circle*) bifurcation point. The SNIC bifurcation ensures that the Ω_1_ oscillator is robustly arrested as the latching parameter increases. (Another possible scenario is that Ω_1_ undergoes a Hopf bifurcation between oscillations and a latched state, as shown in Fig. S1. In this case, there are two different latched states with intermediate and low A_1_ activity.)

Lastly, we study the effects of increasing the checkpoint signal. Because *C*_1_ increases the rate of degradation of I_1_, as *C*_1_ increases, the activation of I_1_ by A_1_ becomes more difficult; hence, the shift of the I_1_ nullcline to higher values of A_1_ (Fig. 3F). This shift generates a new stable steady state; i.e., a *checkpoint*. Unlike the latched state, A_1_ activity is high when the checkpoint is active (Fig. 3F). The bifurcation diagram, A_1_ versus C_1_ (Fig. 3G), shows that increasing the checkpoint signal causes the system’s oscillation to be arrested at a SNIC bifurcation. We note that, from the perspective of the Ω_1_ system, the latched and checkpoint states are ‘symmetric’ with respect to the oscillation. The only notable difference between them is that A_1_ is low in the latched state, but high in the checkpoint state (Fig. 3H).

#### The coupled-oscillator system Ω_1_Ω_2_

We create a coupled-oscillator system by duplicating the Ω_1_ oscillator as Ω_2_ with dynamic variables A_2_ and I_2_, and coupling them (Fig. 4A) by assuming that A_1_ promotes A_2_ degradation in the same way that A_2_ regulates A_1_ in Eq. (1a). Time-course simulations confirm that the components of Ω_1_ and Ω_2_ oscillate out of phase (Fig. 4B), comparable to the out-of-phase oscillations of the S- and M-modules of the cell cycle. The oscillatory regime is surrounded by latched and checkpoint states in case of Ω_1_Ω_2_ coupled oscillators as well (Fig. 4C).

**Figure 4.**
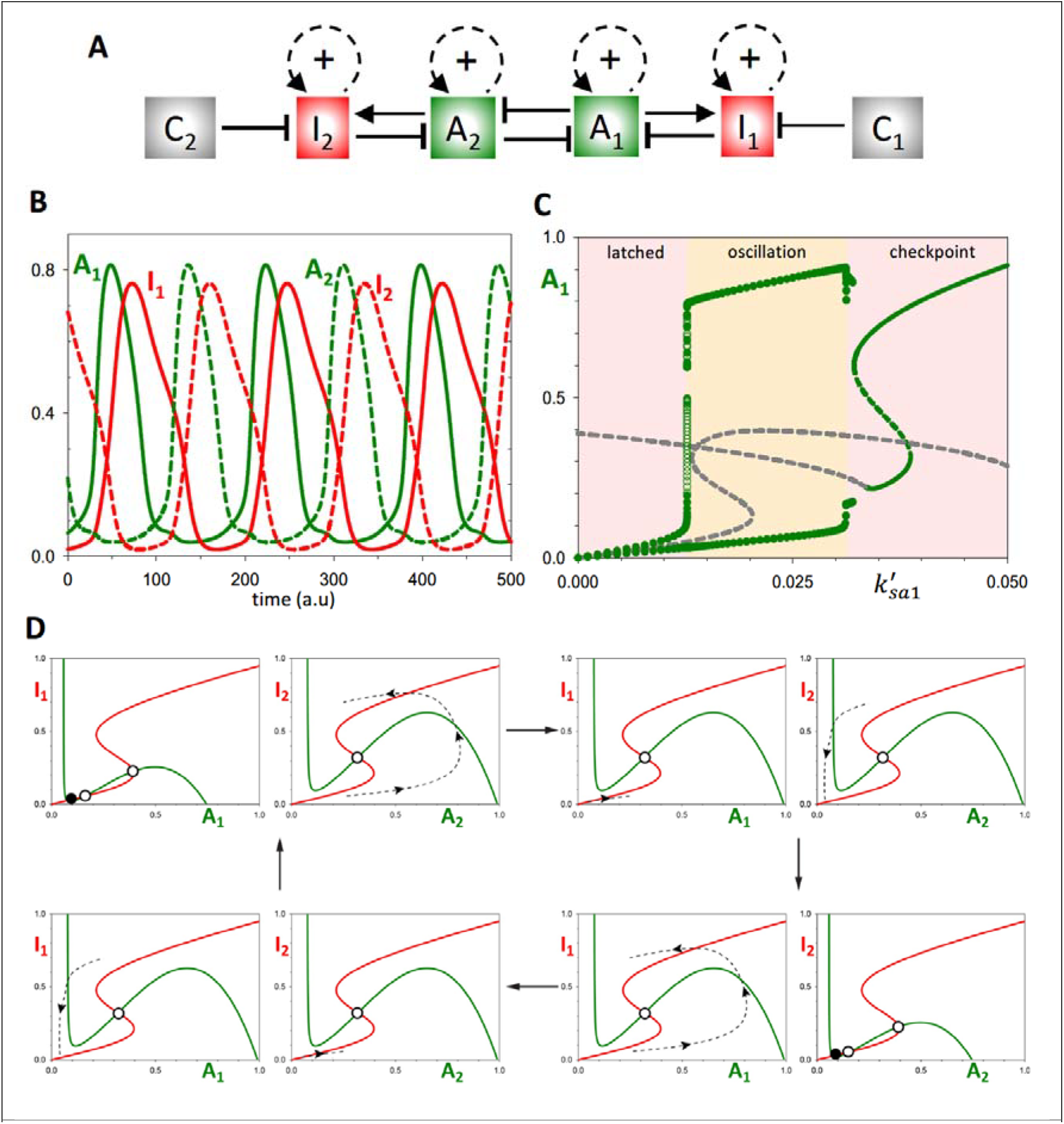
Mutual inhibition of Ω_1_ and Ω_2_ results in out-of-phase oscillations. **(A)** Influence diagram of the Ω_1_Ω_2_ double**-**oscillator network. **(B)** Time course simulation showing the strict alternation of Ω_1_ and Ω_2_ components. **(C)** Bifurcation diagram, *A*_1_ versus basal synthesis rate, *k’*_*sa1*_. **(D)** Phase plane views of the state of the Ω_1_ and Ω_2_ systems at different time points in the coupled oscillation. Notice that, when the level of either activator exceeds ∼0.2 (see Fig. 3E), the other oscillator is latched. When the level of the activator in play falls below this threshold, the other activator unlatches and begins to accumulate.

To understand the mechanism that enables this sequential, unidirectional triggering of the two oscillators, we consider the phase plane of each individual oscillator at different points in the overall cycle (Fig. 4C). As a starting point, we suppose that *A*_2_ is large enough to put Ω_1_ in the latched state with *A*_1_ small, ensuring that Ω_2_ is unlatched and able to oscillate freely.

Consequently, *A*_2_ increases, followed by *I*_2_, followed by an abrupt drop in *A*_2_. As the repression on A_1_ is relieved, the Ω_1_ oscillator is unlatched, *A*_1_ increases, *A*_2_ drops further, and Ω_2_ is latched. Finally, the activity of A_1_ falls, in consequence of negative feedback from I_1_, which unlatches and Ω_2_, allowing *A*_2_ to increase and complete the cycle.

Within this framework, it is inevitable that an endocycle arises when a perturbation arrests one of the oscillators in the latched position, allowing the other to cycle periodically. For instance, suppose the suppression of Ω_1_ by Ω_2_ increases; say, by increasing *k*_*da1*_ (the second-order rate constant for A_2_-dependent degradation of A_1_). Figure 5B illustrates the Ω_2_ endocycle when *k*_*da1*_ = 0.7. Bifurcation diagrams (Fig. 5C,D) reveal that these endocycles arise via a global bifurcation at *k*_*da1*_ ≈ 0.57, where the amplitude of Ω_1_ oscillations drops precipitously, while the Ω_2_ oscillator presses on uninterrupted. An Ω_2_ endocycle could alternatively be generated by silencing the Ω_1_ oscillator internally, e.g., by decreasing the rate of A_1_ accumulation.

**Figure 5.**
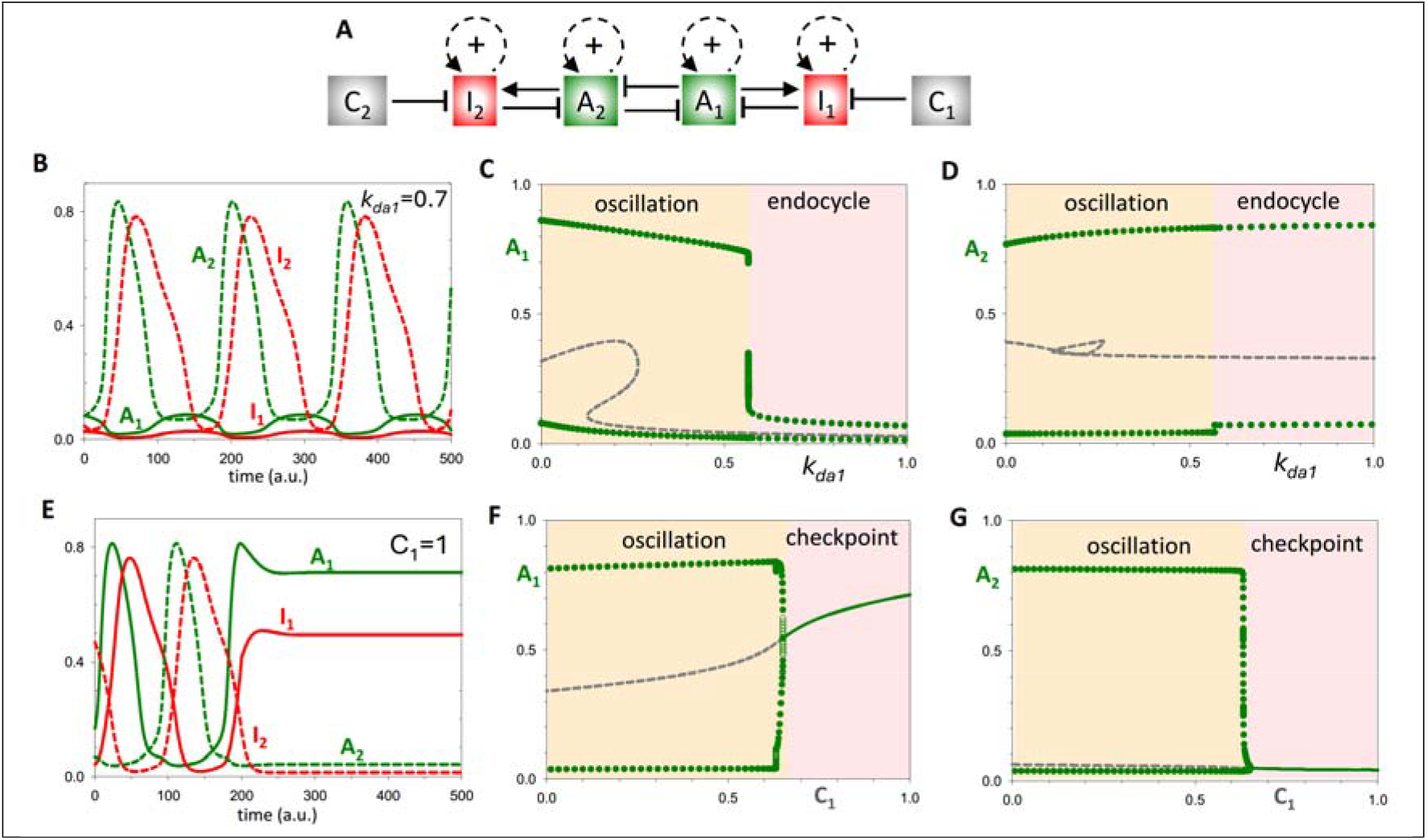
The response of the Ω_1_Ω_2_ double-oscillator to perturbations. **(A)** Influence diagram of the Ω_1_Ω_2_ double**-**oscillator network. **(B)** Time-course simulation after increasing the strength of *A*_2_ inhibition on *A*_1_ (*k*_*da1*_=0.25 → 0.7), which leads to an abrupt transition to endo-oscillations. **(C**,**D)** Bifurcation diagrams illustrating the effects of *k*_*da1*_ on the amplitude of the oscillations. While *k*_*da1*_>0.57 completely suppresses the inherent oscillation of Ω_1_, it has little effect the Ω_2_ oscillation. **(E)** Time-course simulation after increasing the checkpoint parameter *C*_1_=0 → 1, which arrests both oscillators. **(F**,**G)** Bifurcation diagrams show an abrupt transition between oscillatory and checkpoint regimes. Ω_1_ arrests with high activity of *A*_1_, while Ω_2_ arrests in its latched state.

Increasing the checkpoint signal (C_1_) on Ω_1_ arrests both oscillators, as it should (Fig. 5E). By raising the value of *C*_1_, the checkpoint prevents the accumulation of I_1_, such that A_1_ cannot be down-regulated. Consequently, Ω_1_ arrests with a high level of A_1_ (its checkpoint state) and Ω_2_ arrests in its latched state. The bifurcation diagrams, A_1_ and A_2_ versus C_1_ (Fig. 5F,G), show that both oscillators are suppressed at a Hopf bifurcation.

#### The latching-gate perspective

The latching-gate hypothesis frames cell cycle control as a bistable switch between a G1-state of high APC/C:Cdh1 activity and an S/G2/M-state of high Cdk1:CycB activity [27]. The G1/S transition is driven by a ‘helper’ protein (Cdk2:CycE) and the M/G1 transition by a different helper protein (APC/C:Cdc20). A similar view of the double-oscillator model is afforded by the bifurcation diagrams for A_1_ in dependence on I_1_ and I_2_ as helper molecules (Fig. 6B). As expected, the mutual antagonism between A_1_ and A_2_ gives rise to a bistable switch: when (*I*_1_,*I*_2_) ≈ (0,0), there are two alternative, stable steady states: at (*A*_1_,*A*_2_) ≈ (1,0) and at (*A*_1_,*A*_2_) ≈ (0,1). Notice that increasing I_1_ flips A_1_ OFF, but then the mutual antagonism locks Ω_1_ in its OFF state, A_1_≈I_1_≈0 and A_2_≈1. The only way to get A_1_ back to its ON state is by increasing I_2_. By symmetry, the same argument holds for flipping A_2_ ON and OFF. In this manner, the double oscillator executes the full oscillation simulated in Fig. 4B. Notice that, although the mutual antagonism between A_1_ and A_2_ permits a stable steady state at (*A*_1_,*A*_2_) ≈ (0.5,0.5), the cell cycle trajectory never visits this state.

**Figure 6.**
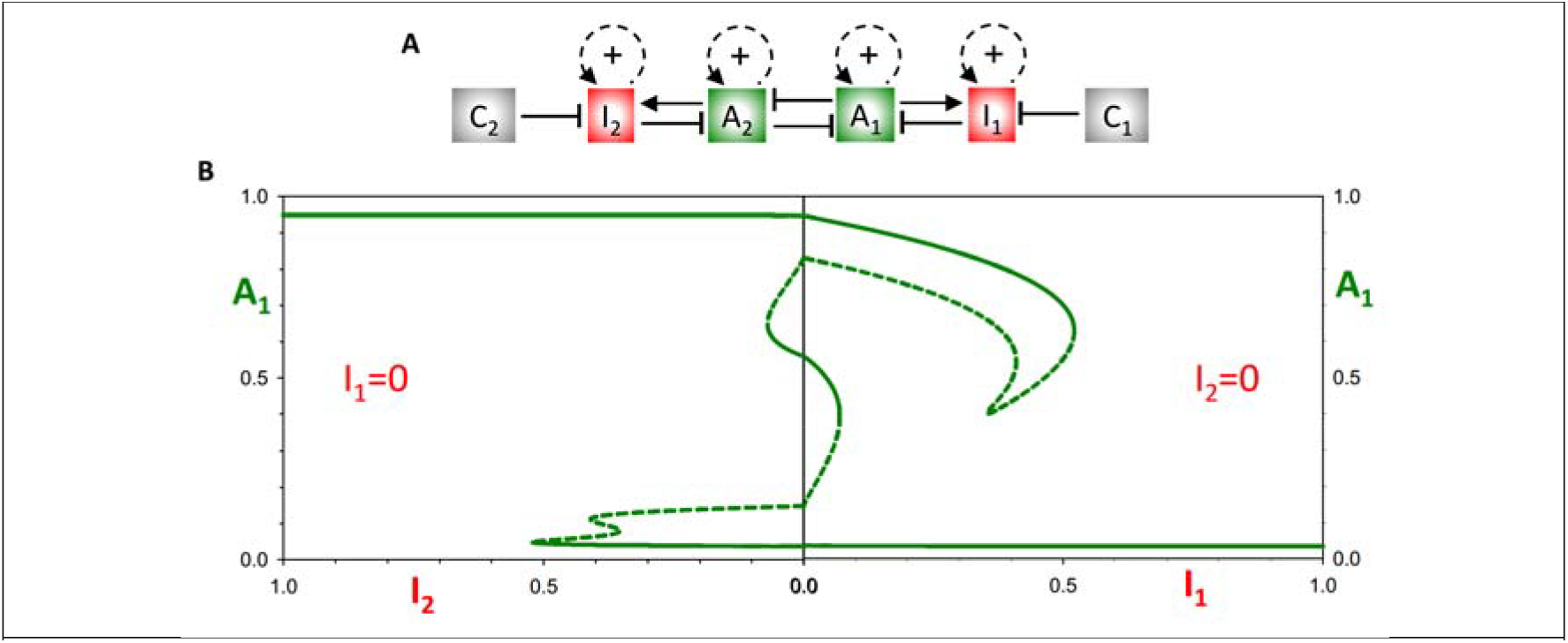
The latching-gate perspective. **(A)** Influence diagram of the Ω_1_Ω_2_ double**-**oscillator network. **(B)** Bifurcation diagrams. (Right panel) *A*_1_ vs I_1_ for I_2_=0. As I_1_ increases, *A*_1_ flips irreversibly from ON to OFF. (Left panel) *A*_1_ vs I_2_ for I_1_=0. As I_2_ increases, *A*_1_ flips irreversibly from OFF to ON. When *A*_1_ is ON, *A*_2_ is OFF, and vice versa, because of the antagonism between *A*_1_ and *A*_2_.

### A mechanistically realistic model of coupled S- and M-oscillators

In this subsection, we return to the mechanistically realistic model of the mammalian cell cycle control network in Fig. 2. As discussed previously, the mechanism consists of two modules for S-phase control and M-phase control, and each module has the features of a doubly amplified, negative feedback loop. Furthermore, the two modules are linked by robust mutual antagonism between APC/C:Cdh1 and Cdk1:CycB. In contrast to the analysis in our published paper [27], where the global dynamics of the system were probed leaving the antagonism between APC/C:Cdh1 and Cdk1:CycB intact, here we begin by breaking the interaction between APC/C:Cdh1 and Cdk1:CycB, in order to analyse the properties of the uncoupled modules and gain understanding of how they influence each other.

To analyse the S-module, we redefine the activity of Cdk1:CycB as a parameter, instead of a dynamic variable (Fig. 7A). Consequently, any effect of the S-module on the M-module is eliminated, as CycA:Cdk1 and APC/C:Cdh1 can no longer influence Cdk1:CycB. Likewise, the level and activity of all the components in the mitotic module remain constant. When Cdk1:CycB activity is zero, the activities of APC/C:Cdc20 and Gwl-kinase are low, while those of B55:PP2A and Wee1 are high. Hence, we can analyse the dynamical properties of the S-module in isolation. To this end, we analyse our system by ‘pseudo-phase plane’ methods, as explained in the Materials & Methods.

**Figure 7.**
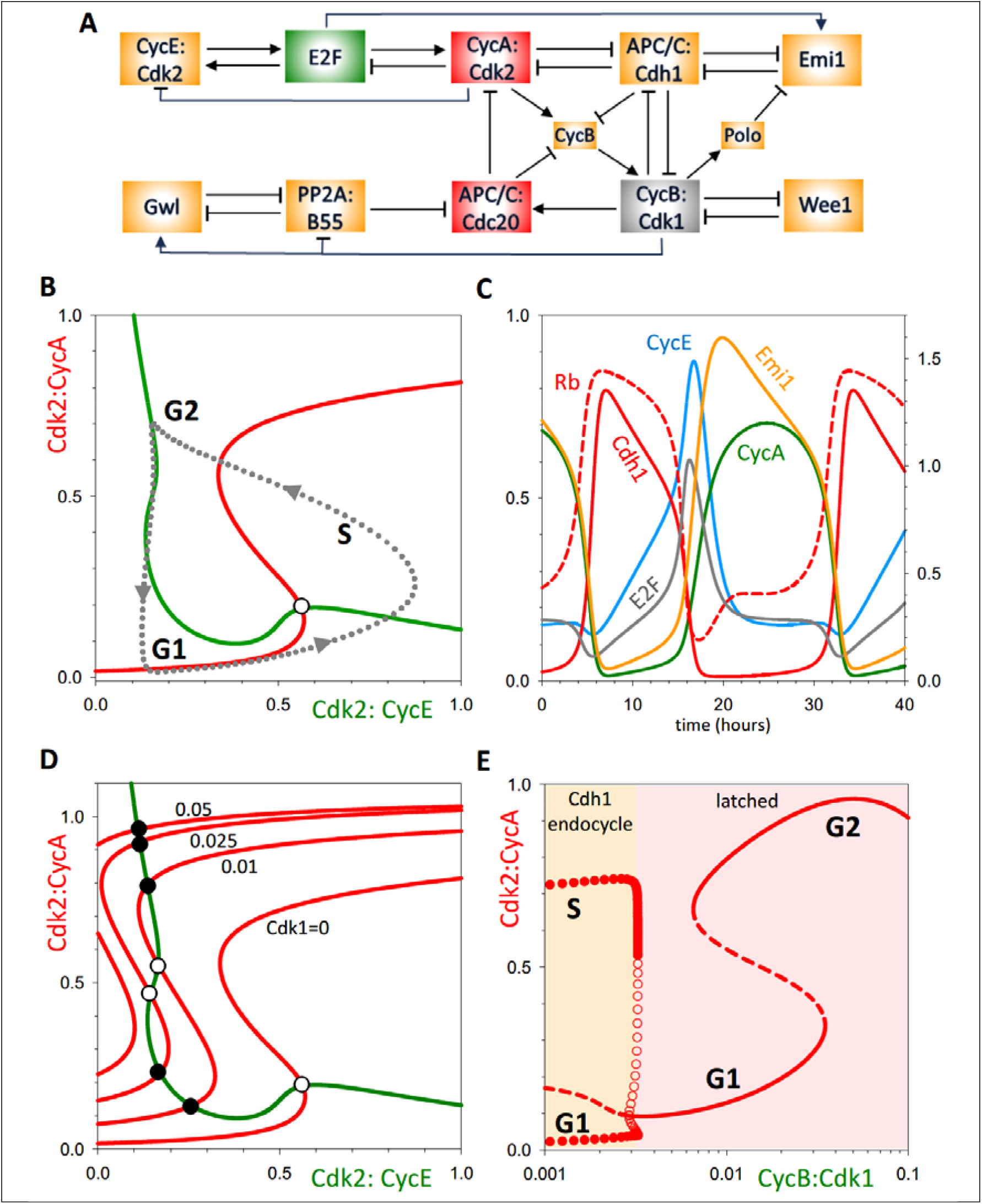
The dynamical properties of the S-phase (Cdh1) oscillator. **(A)** Influence diagram of the system, highlighting the conversion of Cdk1:CycB from a dynamical variable to a parameter (grey box). **(B)** Pseudo-phase plane, Cdk2:CycA versus Cdk2:CycE, exhibits the characteristic features of a doubly amplified negative feedback oscillator. The unstable steady state (o) is surrounded by a stable limit cycle (dashed line is a projection of the time course simulation in panel C). **(C)** Time course simulation. **(D)** Pseudo-phase plane showing the emergence of a stable steady state of Cdk2:CycA activity, as the value of the Cdk1:CycB parameter increases. The stable steady state corresponds to arrest of the S-phase module. **(E)** Bifurcation diagram (Cdk2:CycA activity versus Cdk1:CycB parameter value) showing that a Cdh1-endocycle (endoreplication) emerges abruptly, through a subcritical Hopf bifurcation, when Cdk1:CycB activity falls below a threshold ∼0.0035. In addition, G1- and G2-arrested states are observed at higher values of Cdk1:CycB.

In Fig. 7B, we plot pseudo-nullclines in the (Cdk2:CycE, Cdk2:CycA) phase plane. Cdk2:CycA is the inhibitor of the S-module oscillator. We use Cdk2:CycE as the ‘activator’ of this module because we assume pseudo-steady state kinetics for the binding/dissociation of the Rb:E2F complex, which results in an algebraic equation for E2F as a function of Cdk2:CycE. The Cdk2:CycE nullcline is backwards N-shaped, and the Cdk2:CycA nullcline is S-shaped; analogous to the Ω_1_ oscillator in Fig. 3B. The intersection of the nullclines is an unstable steady state, surrounded by a stable limit cycle. The full time-course of oscillations of the S-phase module (in isolation) is shown on Fig. 7C. As the switching on and off of APC/C:Cdh1 activity is essential to these oscillations, we refer to the S-phase module as a ‘Cdh1-endocycle.’ Because Cdk2:CycA is oscillating with negligible Cdk1:CycB activity, these oscillations constitute endoreplication cycles (repeated rounds of DNA synthesis without mitosis).

In order to understand the coupling between the S-phase and M-phase modules, we now show how increasing the Cdk1:CycB parameter affects the Cdh1-oscillator (Fig.7D, E). Since Cdk1:CycB regulates both the Activator (E2F) and Inhibitor (CycA) of the negative feedback loop, both nulllclines are affected as the value of Cdk1:CycB is increased. Nevertheless, at small Cdk1:CycB values the most relevant changes occur on the Cdk2:CycA-nullcline. Notice how increasing Cdk1 shifts the S-shaped nullcline to the left, making the system bistable, thereby converting the unstable steady state into a stable one, as indicated by the filled circle (Fig.7D). At higher activity of Cdk1:CycB the lower steady states disappear through a saddle-node bifurcation and only the upper stable steady remains. A more detailed picture about the effect of Cdk1:CycB on the Cdh1-oscillator is given by the bifurcation diagram in Fig.7E, which shows that the large amplitude CycA oscillations are terminated by a sub-critical Hopf-bifurcation before the system enters into the bistable regime. For Cdk1 activity >0.04, the system leaves the bistable regime, and only a high CycA-activity (G2) state remains. This dynamic behaviour resembles the situation we saw with our toy-model (Suppl. Fig. S1).

A similar approach can be taken to analyse the mitotic module. For this, we turn the activity of APC/C:Cdh1 into a parameter (indicated by the grey rectangle in Fig.8A), which eliminates its regulation by Cdk2:CycA and Cdk1:CycB. Setting APC/C:Cdh1 to zero (i.e. the system cannot be arrested in G1) and plotting the pseudo-phase plane for Cdk1:CycB and APC/C:Cdc20 (Fig. 8B) reveals nullclines consistent with the mitotic module being an activator-inhibitor-double-amplified negative feedback oscillator. As expected, this system displays limit cycle behaviour (Fig. 8C). Increasing the activity of APC/C:Cdh1 arrests these oscillations through a SNIC (Saddle-node on Infinite Circle) bifurcation (Fig. 8D,E). Again, this is consistent with the hypothesis that the mutual antagonism of the two modules is at the heart of their alternating oscillations. Nevertheless, it must be noted that by setting APC/C:Cdh1 constant, the mitotic modules still receives fluctuating inputs from Cdk2:CycA in the S-phase module. This is caused by APC/C:Cdc20-dependent degradation of CycA, while Cdk2:CycA is responsible for activation of CycB synthesis. To show that the mitotic control module can indeed behave as an autonomous oscillator, we also convert Cdk2:CycA to a parameter. Time course and phase plane analysis reveals the expected properties of the double amplified negative feedback oscillator are still present (Suppl. Fig. S2).

**Figure 8.**
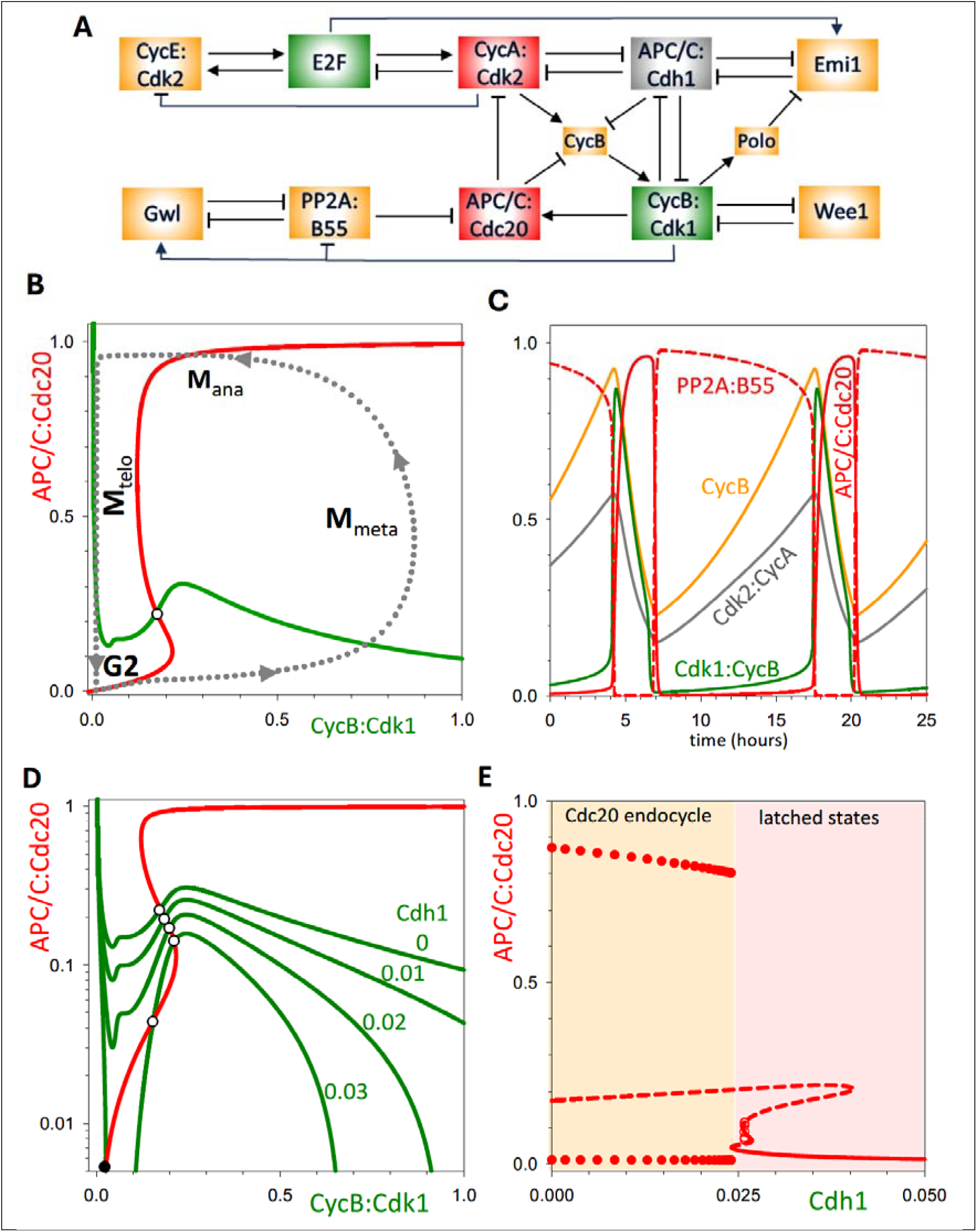
The dynamical properties of the S-phase (Cdh1) oscillator. **(A)** Influence diagram of the system, highlighting the conversion of Cdk1:CycB from a dynamical variable to a parameter (grey box). **(B)** Pseudo-phase plane, Cdk2:CycA versus Cdk2:CycE, exhibits the characteristic features of a doubly amplified negative feedback oscillator. A stable limit cycle (dashed line is a projection of the time course simulation in panel C). **(C)** Time course simulation. **(D)** Pseudo-phase plane showing the emergence of a stable steady state of APC/C:Cdc20 activity, as the value of the APC/C:Cdh1 parameter increases. The stable steady state corresponds to arrest of the M-module. **(E)** Bifurcation diagram (APC/C:Cdc20 activity versus Cdh1 parameter value) showing that a Cdc20-endocycle emerges abruptly, through a SNIC bifurcation, when APC/C:Cdh1 activity falls below a threshold ∼0.025.

## Conclusion

The current work builds on a previously published model of the cell cycle that highlighted the ability of mammalian cells to transition between mitotic cycles and endocycles upon specific perturbations of mitotic kinase activity [27]. Our investigation shows that the basis for these phenomena rests on a mechanism of two activator/inhibitor-amplified negative feedback oscillators that are mutually antagonistic. Using bottom-up modelling combined with phase plane and bifurcation analysis, we have shown that a toy model, consisting of a pair of mutually inhibiting, second order, nonlinear oscillators can reproduce the major <u>qualitative</u> features of mitotic cycles and endocycles; and that a more realistic model, consisting of 14 nonlinear ODEs describing the CDK control system in mammalian cells, recapitulates all the qualitative features of the toy model. Furthermore, the toy model, being a faithful mimic of the real control system, suggests a novel dynamical metaphor for cell cycle control.

### Newton’s cradle

We propose that the alternating oscillation of S- and M-modules (and Ω_1_Ω_2_ oscillators) is analogous to the dynamics of Newton’s cradle (Fig. 9A), a mechanical device used to illustrate the conservation of momentum and other aspects of Newtonian physics. When one of the outer pendula is set in motion, the device executes an alternating sequence of half-swings of both outer pendula. Clearly, the two outer pendula can be associated with the S- and M-modules (Ω_1_Ω_2_ oscillators) of the cell cycle control system. This parallel can be illustrated by plotting pseudo-phase planes for the individual Ω_1_ and Ω_2_ oscillators (Fig. 9B). The only steady state showing up on each of these phase planes corresponds to the latched state of both oscillators, where both activator and inhibitor levels are low. In Newton’s cradle, these steady states correspond to low potential energy positions of the outer spheres, where they are in contact with the other stationary spheres. Once an outer sphere reaches the low point of its oscillation, its momentum is transferred to the other outer sphere. In case of the double-oscillator, reaching one of these latched states causes a release of the other activator from inhibition and drives the system to the other pseudo-phase plane (Fig. 9B). In this way, the motion of Newton’s cradle resembles the alternation of S phase and M phase during the mitotic cycle of eukaryotic cells. As with endocycles, when the oscillators’ coupling is perturbed by blocking the swinging of an outer sphere, the other outermost pendulum oscillates independently as a consequence of elastic collisions with the stationary inner spheres (Fig. S3A,B).

**Figure 9.**
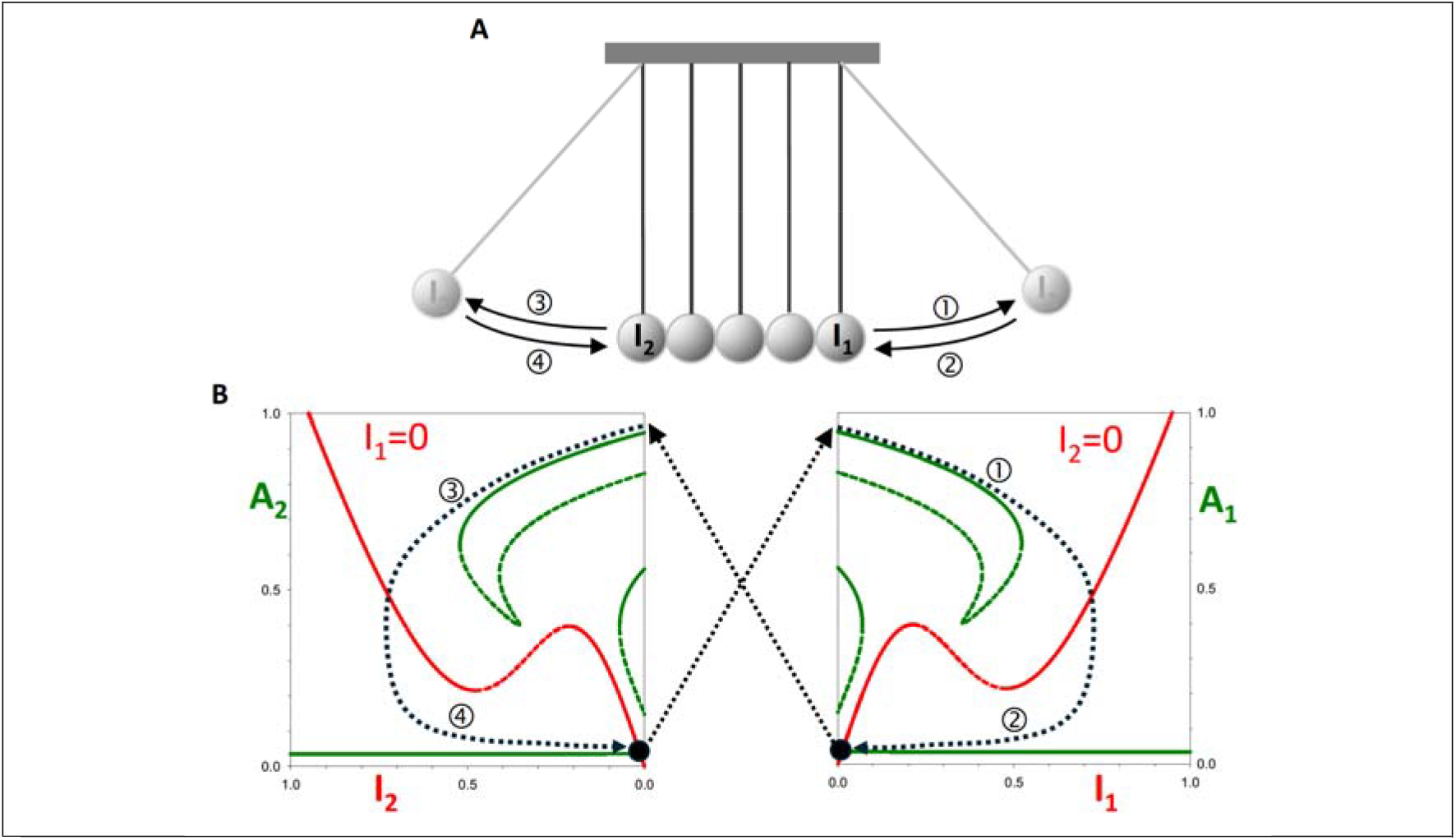
The Newton’s cradle analogy to the double oscillator system. **(A)** Newton’s cradle (see <u>youtube video</u>). **(B)** Pseudo-phase planes for A_1_-I_1_ (right panel) and for A_2_-I_2_ (left panel), and schematic cell cycle trajectory (dotted line). Starting from A_1_≈1 and I_1_≈A_2_≈I_2_≈0, as I_1_ increases A_1_ switches OFF, which allows A_2_ to switch ON, bringing the trajectory to A_2_≈1 and I_2_≈A_1_≈I_1_≈0, completing one-half of the cycle.

Because our double-oscillator model executes limit cycle oscillations in the absence of checkpoints, it is consistent with older clock models (and clock-shop models) of the cell cycle. Furthermore, checkpoints can be seamlessly integrated into our model, easily aligning it with older falling-dominoes and newer latching-gates views of cell cycle controls. In our metaphor of Newton’s cradle, a checkpoint would correspond to catching one of the pendula in a state of high potential energy, thereby preventing oscillations of both outer spheres (Fig. S3C,D). When the checkpoint is released (the sphere is dropped), the normal cycle picks up where it left off. Thus, we suggest that our linked oscillators framework builds on the classical metaphors of cell cycle dynamics and expands them to account elegantly for perturbations leading to endocycles.

## Material and methods

### A doubly amplified negative feedback oscillator

We model a doubly amplified negative feedback oscillator by a pair of nonlinear ordinary differential equations (ODE), Eqs. 1a and 1b, describing the kinetics of Activator-1 (*A*_1_) and Inhibitor-1 (*I*_1_) accumulation and degradation with parameter values in Table1.

**Table 1.** Numerical values of parameters in the two coupled doubly amplified negative feedback oscillators.

| Parameter | Description | Value |
| --- | --- | --- |
| $k'_{sa1}$ and $k'_{sa2}$ | Constitutive synthesis rates of $A_1$ and $A_2$ | 0.015 |
| $k_{sa1}$ and $k_{sa2}$ | Autocatalytic synthesis rates of $A_1$ and $A_2$ | 0.185 |
| $L_{a1}^q$ and $L_{a2}^q$ | Hill coefficients of autocatalytic synthesis of $A_1$ and $A_2$ | 0.01 |
| $q$ | Hill exponent of autocatalytic synthesis of $A_1$ and $A_2$ | 3 |
| $r$ | Inhibition of $A_1$ and $A_2$ synthesis by $I_1$ and $I_2$ | 25 |
| $k'_{da1}$ and $k'_{da2}$ | Constitutive degradation rate constants of $A_1$ and $A_2$ | 0.2 |
| $k_{da1}$ and $k_{da2}$ | Second order rate constants of $A_1$ and $A_2$ degradation | 0.25 |
| $k'_{si1}$ and $k'_{si2}$ | Constitutive synthesis rates of $I_1$ and $I_2$ | 0.05 |
| $k_{si1}$ and $k_{si2}$ | Autocatalytic synthesis rates of $I_1$ and $I_2$ | 0.1 |
| $L_{i1}^p$ and $L_{i2}^p$ | Hill coefficients of autocatalytic synthesis of $I_1$ and $I_2$ | 0.07 |
| $p$ | Hill exponent of autocatalytic synthesis of $I_1$ and $I_2$ | 3 |
| $k'_{di1}$ and $k'_{di2}$ | Constitutive degradation rate constants of $I_1$ and $I_2$ | 0.15 |
| $k_{da1}$ and $k_{da2}$ | Second order rate constants of $I_1$ and $I_2$ degradation | 0.05 |

The model was implemented in freely available software XPPAUT (link). The ‘ode’ file Fig3_FigS1.ode (Supplementary Material) used with XPPAUT to reproduce the time-course simulations (panels C), phase plane diagrams (panels B,D,F) and one-parameter bifurcation diagrams (panels E,G,H) in Figure 3 and Suppl. Fig. S1.

### Two coupled doubly amplified negative feedback oscillators

A second copy (*A*_2_ and *I*_2_) of the doubly amplified NFL was coupled to the first by mutual inhibition, yielding a system of four ODEs:

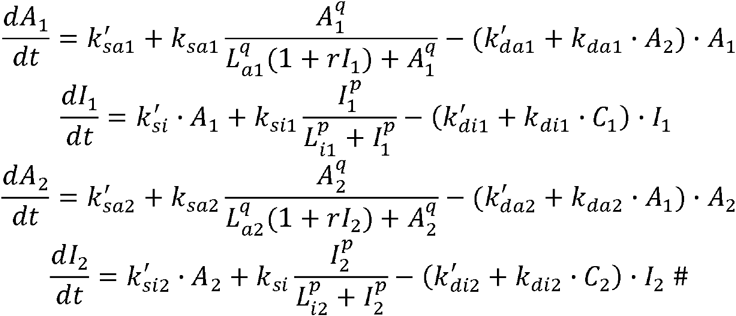

The file Fig4B_Fig5.ode can be used to reproduce the time-course simulations on Fig. 4B and Fig. 5B,E and the one-parameter bifurcation diagrams on Fig. 5C,D,F,G.

Pseudo-nullclines are calculated using the capacity of XPPAUT to plot one-parameter bifurcation diagrams for a reduced set of ODEs. First, the dynamic variable chosen as the bifurcation parameter is removed from the system of ODEs and treated as a parameter. Next, for the *A*_1_*-I*_1_ pseudo-phase plane, *I*_1_ is also converted to a parameter (and similarly for the *A*_2_*-I*_2_ pseudo-phase plane) and a one-parameter bifurcation diagram for the *A*_1_ steady state as a function of *I*_1_ is calculated with the remaining ODEs to plot the pseudo-nullcline for *dA*_1_/*dt* = 0. To calculate the pseudo-nullcline for *dI*_1_/*dt* = 0, the roles of *A*_1_ and *I*_1_ are reversed. The file Fig6_Fig9B.ode calculates the one-parameter bifurcation diagram (i.e., the pseudo-nullcline) of *A*_1_ as a function of *I*_1_.

### A realistic model for the human cell cycle

We used our recently published model for the human cell cycle (Dragoi et al., 2024) with only slight changes to four parameter values. In the file Fig7.ode, both CycB and Cdk1 are converted to parameters, so that it can be used to simulate the oscillatory time-course of endoreplication cycles (Fig. 7C). Fig7.ode also supports calculation of the Cdk2:CycA bifurcation diagram (Fig. 7E) after setting the initial value of the Cdk1 bifurcation parameter to 0.1 in the parameter list. The Cdk2:CycE and Cdk2:CycA pseudo-nullclines in Fig. 7B,D phase planes can be calculated by overlaying two one-parameter bifurcation diagrams. To plot the Cdk2:CycA pseudo-nullcline, CycE is converted to a constant and used as a bifurcation parameter to calculate the CycA steady state values (CycA_ss_). To plot the CycE pseudo-nullcline, CycE is used as the variable and CycA as the bifurcation parameter in a one-parameter bifurcation diagram calculation.

To calculate the Cdc20-endocycles on Fig. 8, Cdh1 was converted from an ODE to parameter in Fig8_FigS2.ode. This file supports calculation of the one-parameter bifurcation diagram of Fig. 8E after setting the initial value of the bifurcation parameter (Cdh1) to 0.05. The limit cycle oscillation at Cdh1=0 can be captured by setting the time of integration to the period of the oscillation (13.35 hours). The pseudo-nullclines for Cdk1:CycB (Cdk1 in the ode file) and for APC/C:Cdc20 are calculated as one-parameter bifurcation diagram after converting Cdc20 or both CycB and Cdk1 into parameters. The file Fig8_FigS2.ode can also be used to calculate all the panels in Suppl. Fig. S2 after setting CycA=0.4 as a parameter using a similar procedure.

## Supporting information

Supplemental ode files

## Acknowledgements

We acknowledge financial support from BBSRC Strategic LoLa grant BB/M00354X/1 to BN.

## Supplementary figures

**Figure S1:**
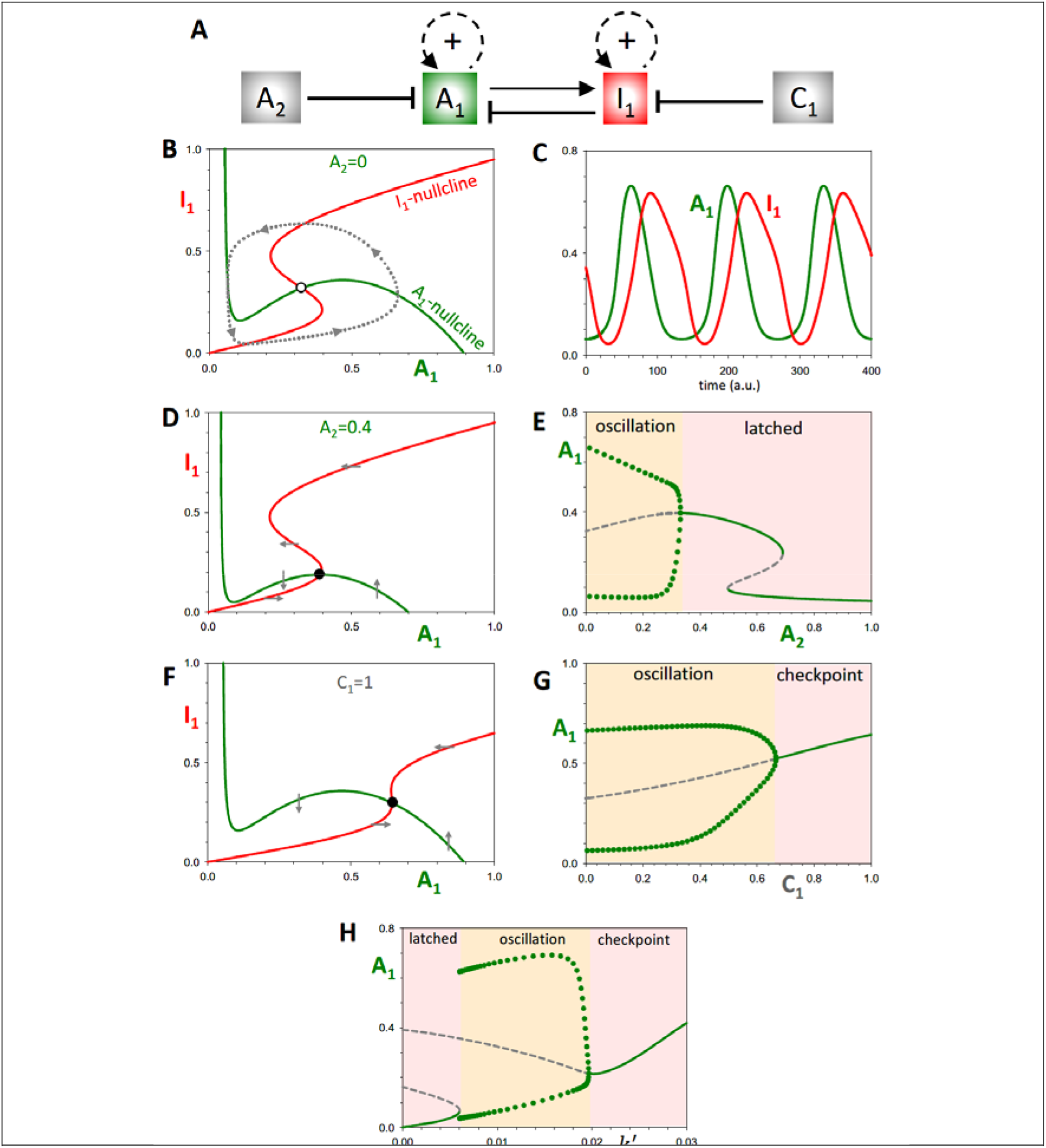
The dynamical properties of the single oscillator, Ω_1_, with an alternative parameter set. **(A)** Influence diagram of the single oscillator network (identical to Fig. 4). **(B)** Phase plane of the *alternative* Ω_1_ system in the basal oscillating state (see Fig3_FigS1.ode file). As in Fig. 4, there is a unique unstable steady state (on the saddles of the two nullclines), surrounded by a limit cycle oscillation. **(C)** Time course simulation. **(D)** Phase plane of the *alternative* Ω_1_ system, when *A*_2_ = 0.4. Notice that, as a result of the shift in the A_1_ nullcline, the unstable steady state is converted to a stable steady state, on the lower stable branch of the Inhibitor nullcline and the upper stable branch of the Activator. **(E)** Bifurcation diagram of A_1_ with respect to the latching parameter, *A*_2_, showing that oscillations are generated at a Hopf bifurcation, as opposed to a SNIC (Fig. 4E). Notice this is similar to the response of the S-phase module to changing Cdk1:CycB activity (Fig. 2E). **(F)** Phase plane for the checkpoint arrested *alternative* Ω_1_ system, assuming *C*_1_ = 1. Observe that the shift in the I_1_ nullcline has resulted in the oscillator unstable steady state being converted to a stable steady state, on the lower stable branch of I_1_ and upper stable branch of A_1_. **(G)** Bifurcation diagram of *A*_1_ with respect to the checkpoint parameter, *C*_1_, showing that oscillations are generated at a Hopf bifurcation. **(H)** Bifurcation diagram of *A*_1_ with respect to the basal synthesis rate, *k*_*sa*_*’*. The plot reveals that checkpoint triggering and latching are symmetric effects from the perspective of the *alternative* Ω_1_.

**Figure S2:**
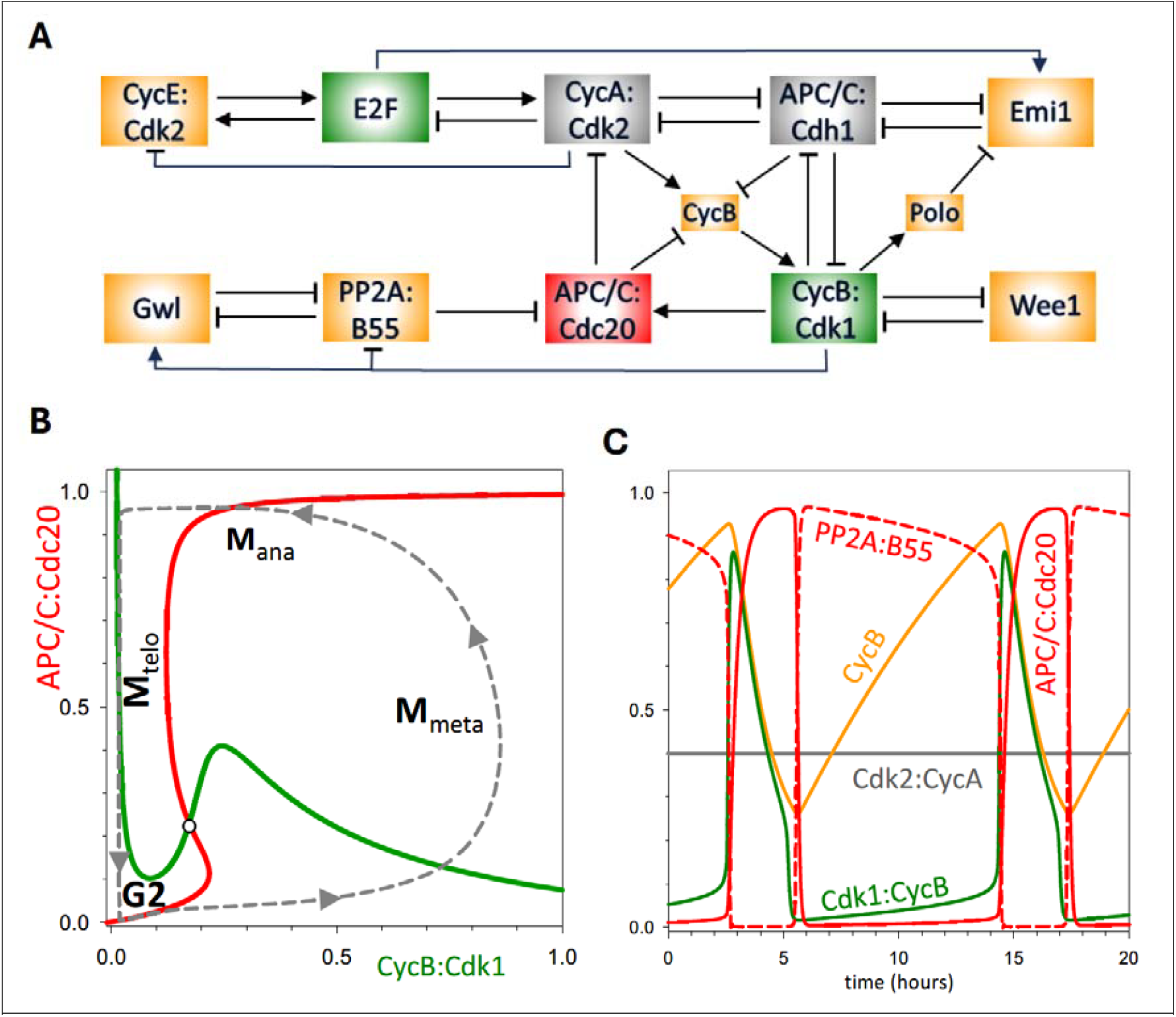
The dynamical properties of the M-module oscillator, for fixed Cdk2:CycA and APC/C:Cdh1 input. **(A)** Influence diagram of the system, highlighting the conversion of APC/C:Cdh1 and Cdk2:CycA from dynamical variables to parameters (grey boxes). **(B)** Phase plane plotting APC/C:Cdc20 against Cdk1:CycB, assuming Cdh1 = 0, CycA = 0.4, showing that the G2/M has intrinsic oscillatory activity in the absence of any dynamical input from the S-phase module. **(C)** The corresponding time course simulation.

**Figure S3:**
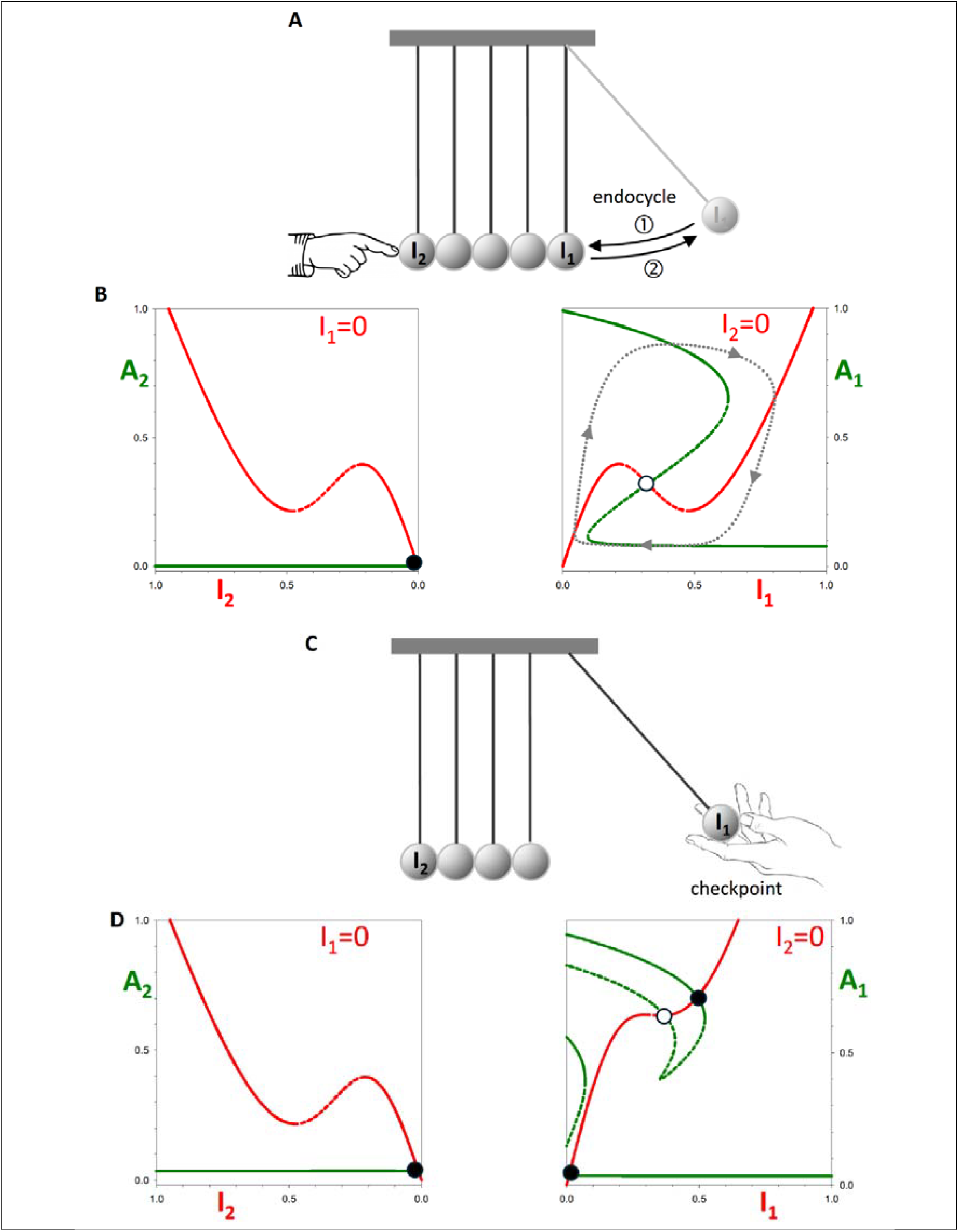
Newton’s cradle analogy to double oscillator perturbations. **(A)** Outline of Newton’s cradle dynamics when the movement of one of the outer spheres is prevented by an external counteracting force. The free-running pendulum continues to swing as a consequence of elastic collisions with the stationary spheres. **(B)** Ω_1_ endocycle pseudo-phase planes, with *k*_*sa2*_*’*=0, showing that the A_2_-I_2_ system is permanently latched, while the A_1_-I_1_ system displays the dynamical properties of a free-running, double-amplified negative feedback oscillator. **(C)** Preventing the return of an oscillating sphere to the equilibrium position prevents the system’s oscillation until the sphere is released. **(D)** Checkpoint pseudo-phase planes, with *C*_1_=1, showing that the generation of a high A_1_ stable steady state prevents oscillations by keeping the A_2_-I_2_ system in the latched position permanently.

