## Supplemental ode files for "Newton’s Cradle: Cell Cycle Regulation by Two Mutually Inhibitory Oscillators"

### Supplementary material

**Fig3_FigS1.ode**

```
# A model for doubly amplified negative feedback oscillator
# A1=Activator-1, I1=Inhibitor-1, A2=Activator-2

# Differential equations
dA1/dt = ksa' + ksa*A1^q/(La^q*(1 + r*I1) + A1^q) - (kda' + kda*A2)*A1
dI1/dt = ksi'*A1 + ksi*I1^p/(Li^p + I1^p) - (kdi' + kdi*C1)*I1

# Parameter values
p A2=0,C1=0,ksa'=0.015,ksa=0.185,kda'=0.2,kda=0.25,La^q=0.01,r=25,q=3
p ksi'=0.05,ksi=0.1,Li^p=0.07,kdi'=0.15,kdi=0.05,p=3

# parameter changes for Fig.S1: ksa'=0.01,ksa=0.19,kda=0.1,La^q=0.1,r=5,q=2

# Fig.3: the period of the oscillation is 154.8 min at A2=0 & ksa'=0.015

# XPPAUT settings
@ nmesh=400,TOTAL=1000,dt=0.5,METH=stiff,XP=A1,YP=I1,XLO=0,XHI=1,YLO=0,YHI=1
@ NTST=150,NMAX=1000000,NPR=10000,DS=-0.01,BOUNDS=200
@ DSMAX=0.001,DSMIN=0.0001,PARMIN=-1,PARMAX=1,AUTOVAR=A1
@ AUTOXMIN=0,AUTOXMAX=0.5,AUTOYMIN=0,AUTOYMAX=1
done
```

**Fig4B_Fig5.ode**

```
# Two doubly amplified negative feedback oscillators with mutual inhibition
# A1=Activator-1, I1=Inhibitor-1, A2=Activator-2, I2=Inhibitor-2

# Differential equations
dA1/dt = ksa' + ksa*A1^q/(La^q*(1 + r*I1) + A1^q) - (kda' + kda1*A2)*A1
dI1/dt = ksi'*A1 + ksi*I1^p/(Li^p + I1^p) - (kdi' + kdi*C1)*I1

dA2/dt = ksa' + ksa*A2^q/(La^q*(1 + r*I2)  + A2^q) - (kda' + kda*A1)*A2
dI2/dt = ksi'*A2 + ksi*I2^p/(Li^p + I2^p) - (kdi' + kdi*C2)*I2

# Parameter values
p kda1=0.25,C1=0,ksa'=0.015,ksa=0.185,kda'=0.2,kda=0.25,La^q=0.01,r=25,q=3
p ksi'=0.05,ksi=0.1,Li^p=0.07,kdi'=0.15,kdi=0.05,p=3,C2=0

# the period of the oscillation is 154.8 (at kda1=0) and 154.9 (at kda1=1)

# Initial conditions
init A1=1,I1=0,A2=0,I2=1

# XPPAUT settings
@ TOTAL=500,dt=0.5,METH=stiff,XLO=0,XHI=500,YLO=0,YHI=1
@ nplot=4,XP=t,YP1=A1,YP2=I1,YP3=A2,YP4=I2
@ NTST=150,NMAX=1000000,NPR=10000,DS=0.01,BOUNDS=200
@ DSMAX=0.001,DSMIN=0.0001,PARMIN=0,PARMAX=1,AUTOVAR=A1
@ AUTOXMIN=0,AUTOXMAX=1,AUTOYMIN=0,AUTOYMAX=1
done
```

**Fig6_Fig9B.ode**

```
# Figure 6 & Figure 9B
# Calculation of pseudo-nullclines for
# two doubly amplified negative feedback oscillators with mutual inhibition
# A1=Activator-1, I1=Inhibitor-1, A2=Activator-2, I2=Inhibitor-2

# Differential equations
dA1/dt = ksa' + ksa*A1^q/(La^q*(1 + r*I1) + A1^q) - (kda' + kda*A2)*A1
# dI1/dt = ksi'*A1 + ksi*I1^p/(Li^p + I1^p) - (kdi' + kdi*C1)*I1

dA2/dt = ksa' + ksa*A2^q/(La^q*(1 + r*I2)  + A2^q) - (kda' + kda*A1)*A2
# dI2/dt = ksi'*A2 + ksi*I2^p/(Li^p + I2^p) - (kdi' + kdi*C2)*I2

# Parameter values
p I1=0,I2=0,ksa'=0.015,ksa=0.185,kda'=0.2,kda=0.25,La^q=0.01,r=25,q=3,C1=0
p ksi'=0.05,ksi=0.1,Li^p=0.07,kdi'=0.15,kdi=0.05,p=3,C2=0

# Initial conditions
init A1=1, A2=0

# XPPAUT settings
@ TOTAL=500,dt=0.5,METH=stiff,XP=t,YP=A1
@ XLO=0,XHI=500,YLO=0,YHI=1
@ NTST=150,NMAX=1000000,NPR=10000,DS=0.001,BOUNDS=200
@ DSMAX=0.001,DSMIN=0.0001,PARMIN=-20,PARMAX=1,AUTOVAR=A1
@ AUTOXMIN=0,AUTOXMAX=1,AUTOYMIN=0,AUTOYMAX=1
done
```

**Fig7.ode**

```
# Model for the human cell cycle and endoreplication cycle
# Differential equations
CycE' = kscyce' + kscyce"*E2F - (kdcyce' + kdcyce"*CycA)*CycE
CycA' = kscyca' + kscyca"*E2F - (kdcyca' + kdcyca"*Cdc20 + kdcyca*Cdh1)*CycA
E2FPt' = kpe2f*(CycA+eps*Cdk1)*(E2FT - E2FPt) - kdpe2f*E2FPt
Rb' = kdprb*(Rbt-Rb)/(Jrb+Rbt-Rb) - kprb*(CycE+CycA+eps*Cdk1)*Rb/(Jrb+Rb)
Emi1' = ksemi1' + ksemi1"*E2F - (kdemi1' + kdemi1"*Cdh1 + kdemi1*Polo)*Emi1
# CycB' = kscycb' + kscycb*CycA - Vdcycb*CycB
# Cdk1' = kscycb' + kscycb*CycA + V25*(CycB - Cdk1) - Vwee*Cdk1 - Vdcycb*Cdk1
Cdh1' = kacdh1*(Cdh1t-Cdh1) - (kicdh1'*CycE+kicdh1"*CycA+kicdh1*eps*Cdk1)*Cdh1
Cdc20' = kacdc20*eps*Cdk1*(1-Cdc20) - kicdc20*PP2AB55*Cdc20
PoloT' = kspolo' + kspolo*CycA - (kdpolo' + kdpolo"*Cdh1)*PoloT
Polo' = (kapolo'*CycA + kapolo"*eps*Cdk1)*(PoloT-Polo)/(Jpolo+PoloT-Polo)-kipolo*Polo/(Jpolo+Polo)
pENSAt' = kGwENSA*pGwl*(ENSAtot - pENSAt) - kcatB55*Complex
pGwl' = (kCdkGwl'*CycA+kCdkGwl*eps*Cdk1)*(Gwtot - pGwl) - (kppx' + kB55Gwl*PP2AB55)*pGwl
PP2AB55' = (kdiss + kcatB55)*Complex-kass*(pENSAt-Complex)*(B55tot-Complex)

# CycB represents the sum of inactive and active Cdk1:CycB dimers, while Cdk1 refers to the active ones only

# Algebraic equations
Rbt = Rbtot/(1 + alpha*SK)
BB2 = Rb + E2FT + Kdrbe2f
Comp2 = (BB2 - sqrt(BB2^2 - 4*Rb*E2FT))/2
E2F = (E2FT-E2FPt)*(E2FT-Comp2)/E2FT
BB1 = Emi1 + Cdh1tot + Kdc1e1
Comp1 = (BB1 - sqrt(BB1^2 - 4*Emi1*Cdh1tot))/2
Cdh1t = Cdh1tot - Comp1
YMEP = GK(kpyme'*CycA+kpyme*eps*Cdk1,kdpyme,Jyme,Jyme)
V25 = k25' + k25*YMEP
Vwee = kwee' + kwee*(1 - YMEP)
Vdcycb = kdcycb' + SAC*kdcycb"*Cdc20 + kdcycb*Cdh1
Complex = B55tot-PP2AB55

# Goldbeter-Koshland function
GB(arg1,arg2,arg3,arg4) = arg2-arg1+arg2*arg3+arg1*arg4
GK(arg1,arg2,arg3,arg4) = 2*arg1*arg4/(GB(arg1,arg2,arg3,arg4)+sqrt(GB(arg1,arg2,arg3,arg4)^2-4*(arg2-
arg1)*arg1*arg4))

# Parameter values
p Cdk1=0, kscyce'=0, kscyce"=1.5, kdcyce'=0.6, kdcyce"=1.5
p kscyca'=0, kscyca"=0.45, kdcyca'=0.045, kdcyca"=0.75, kdcyca=3.75
p Rbtot=1.75, JRb=0.1, kprb=15, kdprb=10.5, alpha=1, SK=1
p E2FT=1, kdpe2f=0.3, kpe2f=1.5, Kdrbe2f=0.001
p ksemi1'=0, ksemi1"=1.5, kdemi1'=0.18, kdemi1"=3, kdemi1=7.5, Kdc1e1=0.0175
p Cdh1tot=1, kacdh1=15, kicdh1'=15, kicdh1"=30, kicdh1=6000
p kscycb'=0, kscycb=0.3, kdcycb'=0.06, kdcycb"=0.75, kdcycb=3.75
p kpyme'=0, kpyme=30, kdpyme=6, Jyme=0.1, k25'=0.45, k25=15, kwee'=0.15, kwee=15
p kicdc20=15, kacdc20=3, eps=1, SAC=1
p kspolo'=0.15, kspolo=0, kdpolo'=0.15, kdpolo"=15
p kapolo'=0, kapolo"=15, kipolo=7.5, Jpolo=0.01
p ENSAtot=4, B55tot=1, kass=7500, kdiss=4.5, kcatB55=15
p kGwENSA=15, kppx'=6, kCdkGwl'=0, kCdkGwl=30, kB55Gwl=60, Gwtot=1

### XppAut SETTINGS
@ Method=stiff, Total=40, Bounds=100, Dt=0.5, tol=1e-5
@ nplot=4,XP=time,YP=Rb,YP2=CycE,YP3=CycA,YP4=Cdh1,Xlo=0,Xhi=40,Ylo=0,Yhi=1
@ NTST=150,NMAX=1000000,NPR=10000,DS=-0.01,BOUNDS=200
@ DSMAX=0.01,DSMIN=0.001,PARMIN=0,PARMAX=2, AUTOVAR=CycA
@ AUTOXMIN=0,AUTOXMAX=0.1,AUTOYMIN=0,AUTOYMAX=1
done
```

**Fig8_FigS2.ode**

```
# Model for the human cell cycle and Cdc20-endocycle
# Differential equations
CycE' = kscyce' + kscyce"*E2F - (kdcyce' + kdcyce"*CycA)*CycE
CycA' = kscyca' + kscyca"*E2F - (kdcyca' + kdcyca"*Cdc20 + kdcyca*Cdh1)*CycA
E2FPt' = kpe2f*(CycA+eps*Cdk1)*(E2FT - E2FPt) - kdpe2f*E2FPt
Rb' = kdprb*(Rbt-Rb)/(Jrb+Rbt-Rb) - kprb*(CycE+CycA+eps*Cdk1)*Rb/(Jrb+Rb)
Emi1' = ksemi1' + ksemi1"*E2F - (kdemi1' + kdemi1"*Cdh1 + kdemi1*Polo)*Emi1
CycB' = kscycb' + kscycb*CycA - Vdcycb*CycB
Cdk1' = kscycb' + kscycb*CycA + V25*(CycB - Cdk1) - Vwee*Cdk1 - Vdcycb*Cdk1
# Cdh1' = kacdh1*(Cdh1t-Cdh1) - (kicdh1'*CycE+kicdh1"*CycA+kicdh1*eps*Cdk1)*Cdh1
Cdc20' = kacdc20*eps*Cdk1*(1-Cdc20) - kicdc20*PP2AB55*Cdc20
PoloT' = kspolo' + kspolo*CycA - (kdpolo' + kdpolo"*Cdh1)*PoloT
Polo' = (kapolo'*CycA + kapolo"*eps*Cdk1)*(PoloT-Polo)/(Jpolo+PoloT-Polo)-kipolo*Polo/(Jpolo+Polo)
pENSAt' = kGwENSA*pGwl*(ENSAtot - pENSAt) - kcatB55*Complex
pGwl' = (kCdkGwl'*CycA+kCdkGwl*eps*Cdk1)*(Gwtot - pGwl) - (kppx' + kB55Gwl*PP2AB55)*pGwl
PP2AB55' = (kdiss + kcatB55)*Complex-kass*(pENSAt-Complex)*(B55tot-Complex)

# CycB represents the sum of inactive and active Cdk1:CycB dimers, while Cdk1 refers to the active ones only

# Algebraic equations
Rbt = Rbtot/(1 + alpha*SK)
BB2 = Rb + E2FT + Kdrbe2f
Comp2 = (BB2 - sqrt(BB2^2 - 4*Rb*E2FT))/2
E2F = (E2FT-E2FPt)*(E2FT-Comp2)/E2FT
BB1 = Emi1 + Cdh1tot + Kdc1e1
Comp1 = (BB1 - sqrt(BB1^2 - 4*Emi1*Cdh1tot))/2
Cdh1t = Cdh1tot - Comp1
YMEP = GK(kpyme'*CycA+kpyme*eps*Cdk1,kdpyme,Jyme,Jyme)
V25 = k25' + k25*YMEP
Vwee = kwee' + kwee*(1 - YMEP)
Vdcycb = kdcycb' + SAC*kdcycb"*Cdc20 + kdcycb*Cdh1
Complex = B55tot-PP2AB55

# Goldbeter-Koshland function
GB(arg1,arg2,arg3,arg4) = arg2-arg1+arg2*arg3+arg1*arg4
GK(arg1,arg2,arg3,arg4) = 2*arg1*arg4/(GB(arg1,arg2,arg3,arg4)+sqrt(GB(arg1,arg2,arg3,arg4)^2-4*(arg2-
arg1)*arg1*arg4))

# Parameter values
p Cdh1=0, kscyce'=0, kscyce"=1.5, kdcyce'=0.6, kdcyce"=1.5
p kscyca'=0, kscyca"=0.45, kdcyca'=0.045, kdcyca"=0.75, kdcyca=3.75
p Rbtot=1.75, JRb=0.1, kprb=15, kdprb=10.5, alpha=1, SK=1
p E2FT=1, kdpe2f=0.3, kpe2f=1.5, Kdrbe2f=0.001
p ksemi1'=0, ksemi1"=1.5, kdemi1'=0.18, kdemi1"=3, kdemi1=7.5, Kdc1e1=0.0175
p Cdh1tot=1, kacdh1=15, kicdh1'=15, kicdh1"=30, kicdh1=6000
p kscycb'=0, kscycb=0.3, kdcycb'=0.06, kdcycb"=0.75, kdcycb=3.75
p kpyme'=0, kpyme=30, kdpyme=6, Jyme=0.1, k25'=0.45, k25=15, kwee'=0.15, kwee=15
p kicdc20=15, kacdc20=3, eps=1, SAC=1
p kspolo'=0.15, kspolo=0, kdpolo'=0.15, kdpolo"=15
p kapolo'=0, kapolo"=15, kipolo=7.5, Jpolo=0.01
p ENSAtot=4, B55tot=1, kass=7500, kdiss=4.5, kcatB55=15
p kGwENSA=15, kppx'=6, kCdkGwl'=0, kCdkGwl=30, kB55Gwl=60, Gwtot=1

### XppAut SETTINGS
@ Method=stiff,Total=25,Bounds=100,Dt=0.1,tol=1e-5,Xplot=time,YPlot=Cdk1,Xlo=0,Xhi=25,Ylo=0,Yhi=1
@ NTST=150,NMAX=1000000,NPR=10000,DS=0.01,BOUNDS=200
@ DSMAX=0.01,DSMIN=0.001,PARMIN=-1,PARMAX=1, AUTOVAR=Cdk1
@ AUTOXMIN=0,AUTOXMAX=0.05,AUTOYMIN=0,AUTOYMAX=1
done
```
